# Parallel tracking by sequencing reveals the impact of restriction modification systems on transfer of an integrative and conjugative element

**DOI:** 10.64898/2026.02.02.703224

**Authors:** Emily B. Gee, Ryan D. Kuster, Dawn M. Klingeman, Joshua K. Michener

## Abstract

Horizontal gene transfer (HGT) is a promising avenue for microbiome engineering that enables DNA delivery to microbes in their native environment. Integrative and conjugative elements (ICEs), a class of genomically-integrated mobile elements, are ideal vectors for this purpose. Developing targeted ICE transfer for controlled microbiome engineering requires a better understanding of ICE mobility in complex communities. However, current methods for tracking horizontal gene transfer are not high throughput. To improve upon current methods, we developed a sequencing-based strategy to track and quantify ICE transfer in complex mixtures that we refer to as ‘ICE-seq.’ This method was able to assess ICE transfer in a synthetic community of 150 recipients simultaneously and in combination with phenotypic assays demonstrated that restriction-modification (RM) systems in the donor and recipient alter ICE transfer rate with subspecies resolution by up to 10,000-fold. We propose that manipulating RM systems can be a strategy to engineer targeted ICE transfer for directed microbiome engineering.

**Importance:** Typical approaches to engineer microbiomes involve adding beneficial strains to existing microbiomes, for example the addition of probiotics to the human gut microbiome or soil amendments to improve plant health or for bioremediation. However, these beneficial bacteria often do not survive for extended periods of time, which limits their overall effectiveness. To circumvent this issue, we can transfer beneficial genes by native mechanisms into existing bacteria that are already in the target environment where they can then perform the beneficial function encoded by these genes. In this work, we developed a technique to track gene transfer into multiple recipients simultaneously. Furthermore, bacteria have innate defense systems to protect against invading DNA that can reduce this gene transfer. Our technique allowed us to identify the impact of these defense systems and their potential function for targeting gene transfer to specific recipients.

## Introduction

Microbiomes such as those associated with the plant rhizosphere or found in the human gut are critically important to the health of the host (1, 2). Many years of co-evolution have created complex interactions that provide benefits to both (3). Modifying microbiomes for beneficial outcomes is a promising approach because of their wide-reaching effect and applicability. However, microbiome engineering attempts are often limited by the establishment of non-native microbes (4). Often, introduced microbes do not establish well and are instead outcompeted by native microbes that are better adapted to the environment (5–8). This establishment barrier limits the long-term effectiveness of probiotics and microbial soil amendments (9).

Horizontal gene transfer is an alternative approach to introduce beneficial genes into a microbiome *in situ* without requiring the establishment of a new microbe. In this way, the introduced “donor” strain only needs to persist long enough to deliver its genetic cargo to native microbes, which can then perform the desired function. There are several types of horizontal gene transfer that can be harnessed for gene delivery including transformation, transduction, and conjugation (10). Transformation, the uptake of external DNA, is generally not a feasible method for targeted microbiome engineering because DNA can degrade in the environment and transformation requires natural competence (11, 12). Transduction, the transfer of DNA by phage, can be more easily targeted because bacteriophages are often very specific to certain microbes (13). However, transduction is limited by host range and stability in the environment, and faces challenges such as acquired bacterial resistance (14, 15). Conjugation is the transfer of DNA between bacterial cells by direct contact, typically by conjugative plasmids. Broad-host-range plasmids can deliver large amounts of genetic cargo, but often lack specificity and stability (16–18). In contrast, integrative and conjugative elements (ICEs) both have a relatively large capacity for genetic cargo and are typically stable over time because they are integrated in the host genome (19). These factors make ICEs ideal vectors for gene delivery.

ICE*Bs*1 is a well-studied ICE native to *Bacillus subtilis*, which integrates site-specifically in a conserved leucine tRNA gene (20). A previous study has shown this ICE’s ability to transfer into several Gram-positive bacteria native to the soil and human gut (21). However, much is still unknown with regards to its transfer rate and breadth, which limits its precision for microbiome engineering. Additionally, its potential for targeted transfer remains unclear. There are several factors that may result in donor-recipient incompatibilities that can restrict ICE transfer. Because conjugation requires direct physical contact, membrane composition and outer membrane proteins are involved in cell attachment and mating pair formation (22). Once in the new cell, ICE integration requires complementary strand synthesis as well as integrase activity in the new host (23, 24). Furthermore, microbial defense mechanisms such as restriction modification (RM) systems or CRISPR systems may also limit horizontal gene transfer (25, 26). Microbial RM systems’ involvement in limiting DNA transformation and protection against phage invasion have been well documented, but their role in conjugation has not been well-explored (27–31).

Some phages, conjugative plasmids, and ICEs encode anti-restriction proteins that can help prevent DNA cleavage in the new host (32, 33). Because conjugation transfers only a single strand of DNA, after complementary strand synthesis has occurred in the new host, conjugated DNA is only hemi-methylated (34). Thus, RM systems are traditionally thought to play little role in conjugation compared to other types of horizontal transfer involving dsDNA (35–38).

While high transfer rates into desired recipients is important for efficient microbiome engineering, limiting transfer into undesired recipients is also important to mitigate off-target effects. Tracking modifications and potential off-target effects can be difficult because of inherently complex dynamics found in native microbiomes (39). Therefore, to engineer microbiomes in a targeted way, sensitive strategies are needed to track these changes. One of the most straightforward strategies to track gene transfer involves encoding an antibiotic resistance gene on a mobile genetic element of interest and using selective culturing to isolate transformed recipients (40). However, these techniques typically must be fine-tuned for each recipient and are not easily applied to mixed communities. A similar approach can be used with fluorescence and cell sorting to identify recipients that have received a fluorescence gene (41). Sequencing-based approaches can also be a powerful tool, because unlike these methods, they do not require live cells. High-throughput chromosome conformation capture (Hi-C) can be used to map conjugation by cross-linking the DNA of mobile genetic elements to their host genome prior to sequencing (42). Environmental transformation sequencing (ET-seq) has been used to track conjugation and electroporation of a plasmid into members of a synthetic microbial community (43). An RNA-based technique, RNA addressable memory (RAM) is another sequencing-based approach to track plasmid conjugation by using a ribozyme to add barcodes to 16s RNA (44). The advantage that these sequencing-based approaches share is their ability to assess multiple recipients at once without relying on tedious phenotypic assays, making them ideal for monitoring DNA exchange in microbiomes.

To better understand the rate and breadth of transfer of ICE*Bs*1 in complex communities, high-throughput methods are necessary. To this end, we developed a sequencing-based technique called ‘ICE-seq’ that allowed us to track and quantify ICE transfer into multiple recipients simultaneously (Figure 1). Furthermore, ICE-seq combined with phenotypic assays allowed us to identify how RM systems can alter ICE transfer efficiency into specific recipients. We suggest RM systems could be used in the future to target ICEs to specific members of the microbiome.

**Figure 1:**
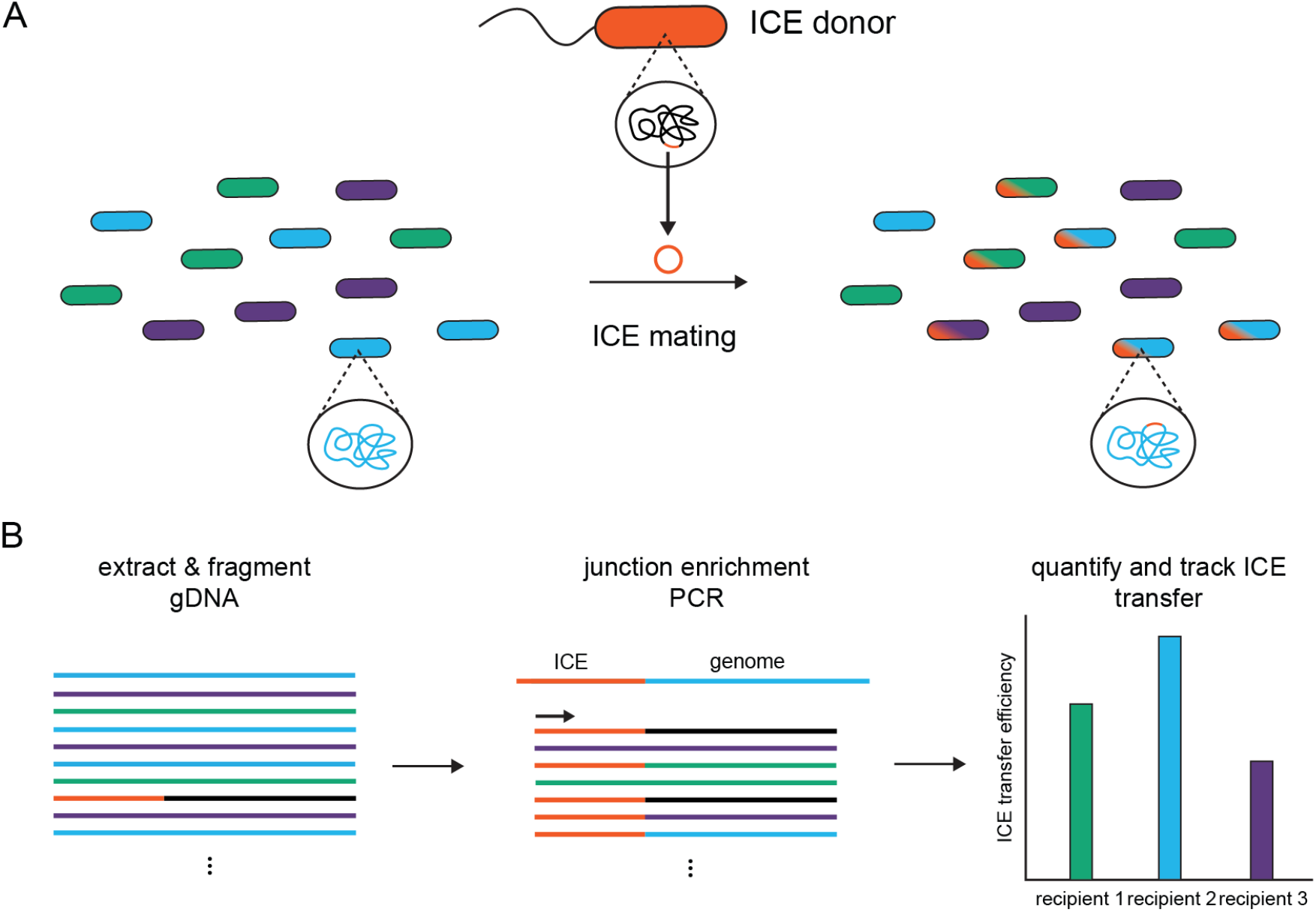
Overview of ICE-seq strategy. **A)** The ICE donor is mixed with a recipient community to allow ICE mating. **B)** Total genomic DNA is extracted from the mixed cells after mating. The DNA is sheared, size-selected, and prepared for NGS sequencing. After adapter ligation, a junction enrichment PCR step amplifies reads at the region of interest and also adds sequencing adapters and indices. By using a primer specific to the region of interest and the other annealing to the sequencing adapter, reads can be amplified from the ICE region into an unknown genomic region. Number of reads matching the transconjugant and donor or recipient are identified and counted to quantify the ICE transfer efficiency.

## Results and Discussion

### ICE-Seq Methods Development

ICE*Bs*1 integrates at an *att* site located in a conserved tRNA gene found in many Gram-positive species. Just outside of this highly conserved region, we identified a region of diversity suitable for differentiating recipient strains by sequencing (Fig. S1). ICE conjugation occurs at low levels under ideal conditions, between 10^-3^ and 10^-8^ transconjugants per donor. Therefore, in nature where transfer rate can be even lower, it would be difficult to detect with metagenomic sequencing and nearly impossible to quantify. Taking inspiration from Tn-seq, we hypothesized that an enrichment step would target reads to the region of interest and enable quantification of ICE transfer even at low frequency in complex mixtures (45, 46).

For ICE-seq, first, the ICE donor was added to a microbial community and allowed to mate. Then, total genomic DNA was extracted from the mixed community without phenotypic selection. DNA fragments at the boundary of the ICE and recipient genome are amplified by junction enrichment PCR (Figure 1). Strategic primer design can target reads that begin in the ICE sequence and end in an unknown region to detect any strain that has integrated the ICE in its genome. A custom script was used to find kmers that are unique to a single recipient and close enough in proximity to the *att* site that they can be captured in a single sequencing read. To ensure diagnostic regions are divergent enough, kmers were required to be greater than a defined Levenshtein distance or the number of edits needed to convert one string into another. The occurrence of these diagnostic kmers in the sequencing data was then counted and recorded (Figure 2A). We selected a kmer size of 31 and an Levenshtein distance of 3 for all analyses.

**Figure 2:**
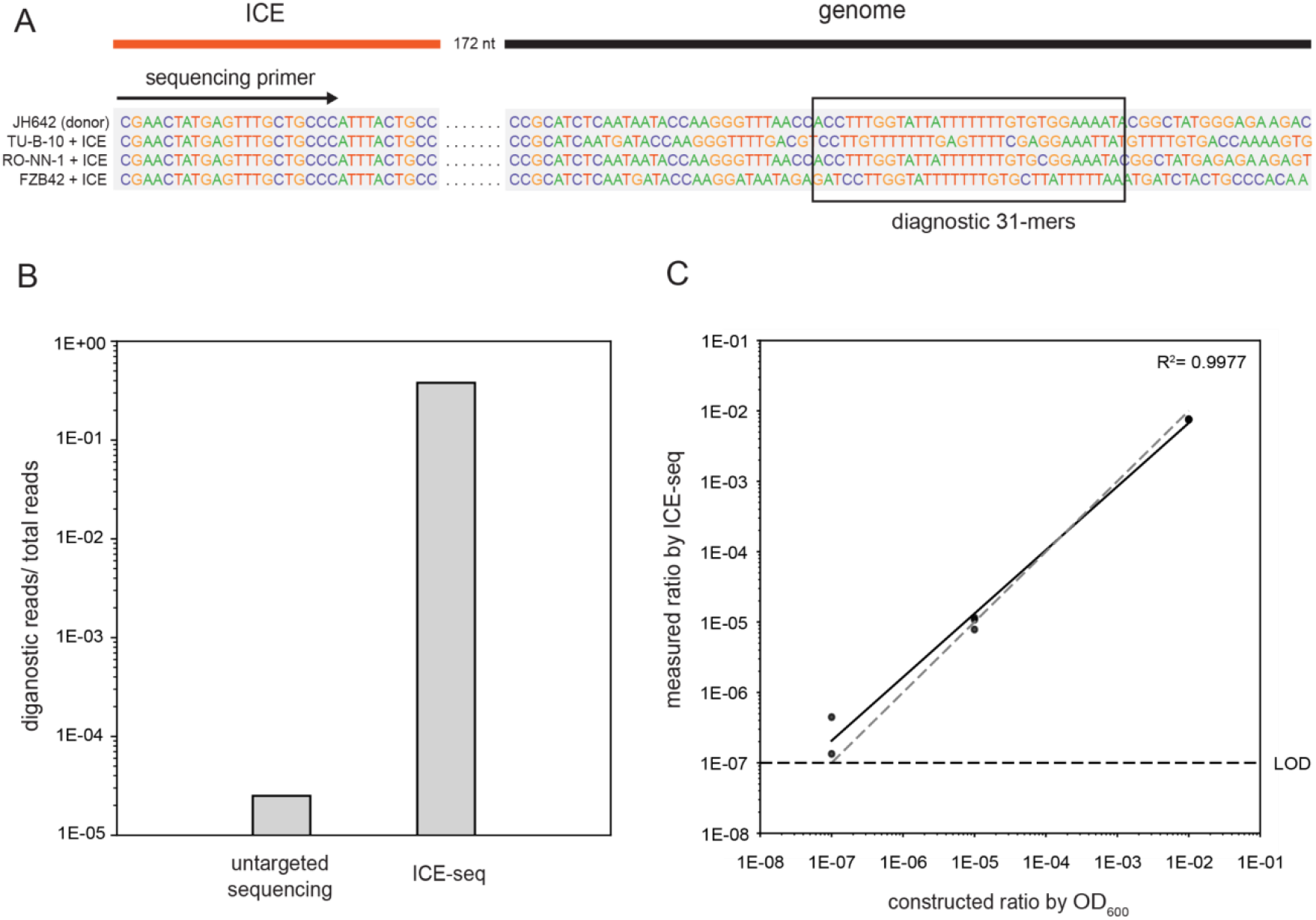
ICE-seq Methods development. **A)** Unique kmers located the same distance from the sequencing primer were identified in the input reference sequences. To quantify ICE transfer, the number of reads containing a specific kmer were counted and used to calculate ICE transfer efficiency. **B)** Untargeted genomic sequencing of a single recipient strain yielded ∼0.0025% of total reads that mapped to the ICE att site. After enrichment of a representative sample using ICE-seq, 37% of reads contain the *att* site. **C)** Known concentrations of isogenic recipient strains containing the ICE (TU-B-10 + ICE/donor = 10-7, RO-NN-1 + ICE/donor = 10^-5^, and FZB42 + ICE/donor = 10^-2^) were mixed by OD_600_ in triplicate and sequenced using ICE-seq. Data was analyzed and plotted by known concentration of strain per donor vs. ratio measured by ICE-seq. Grey dashed line represents x=y, and black line represents log-log linear regression of our data where *R*^2^ = 0.998.

Junction enrichment allowed a large portion of sequencing reads to be found at the region of interest. Compared to whole genome sequencing of a single transconjugant strain where ∼0.0025% of reads contained the *att* site, with ICE-seq around 37% of reads contained the *att* site. (Figure 2B). To test if this method could be used quantitatively, isogenic strains containing the ICE were constructed by conjugation and isolated by selective plating. These were mixed with the donor strain at known concentrations as measured by OD_600_. Genomic DNA was extracted directly from cells after mixing and sequenced with ICE-seq. The ratios measured by ICE-seq were very similar to the constructed ratios, demonstrating that ICE-seq can accurately measure transconjugant frequencies across 5 orders of magnitude. The limit of detection was approximately 10^-7^ using an average sequencing depth of 46 million reads per sample. (Figure 2C, *R*^2^=0.998, standard linear regression).

### ICE-seq tracks and quantifies ICE transfer in mixed populations

To test if ICE-seq can be used to track and and quantify ICE transfer with multiple recipients, we chose five *Bacillus* recipients (*Bacillus spizizenii* TU-B-10, *Bacillus subtilis* subsp. *subtilis* RO-NN-1, *Bacillus amyloliquefaciens* DSM 7, *Bacillus velezensis* FZB42, and *Bacillus velezensis* GB03) that contain the ICE*Bs*1 *att* site and a region of genetic diversity in the surrounding genomic region (Fig. S1). Recipients were mixed with the ICE donor at a one to one ratio by OD_600_ (combined recipients to donor), isolated by vacuum-filtration, and allowed to filter-mate for two hours. Genomic DNA was extracted from the mixed cells and sequenced by ICE-seq. As a proof of principle, individual recipients were also tested in pairwise assays. Recipient strains were mixed with the donor at a one to one ratio by OD_600_ (single recipient to donor) In the pairwise samples, the primer selection was also different (Figure 3A). The primers used in pairwise assays sequence from a unique region in the recipient genome into the *att* site to quantify presence versus absence of the ICE. Additionally, pairwise samples were both sequenced and grown on selective plates to quantify ICE transfer phenotypically (Figure 3B). When measured by sequencing, transconjugant frequencies for pairwise matings and mixed matings were statistically indistinguishable for all recipients (Figure 3B, n = 3, t-test, P<0.05). Recipients differed in the frequency of ICE transfer by approximately two orders of magnitude between 10^-3^ and 10^-5^, suggesting that intraspecific genetic differences quantitatively affect ICE transfer.

**Figure 3:**
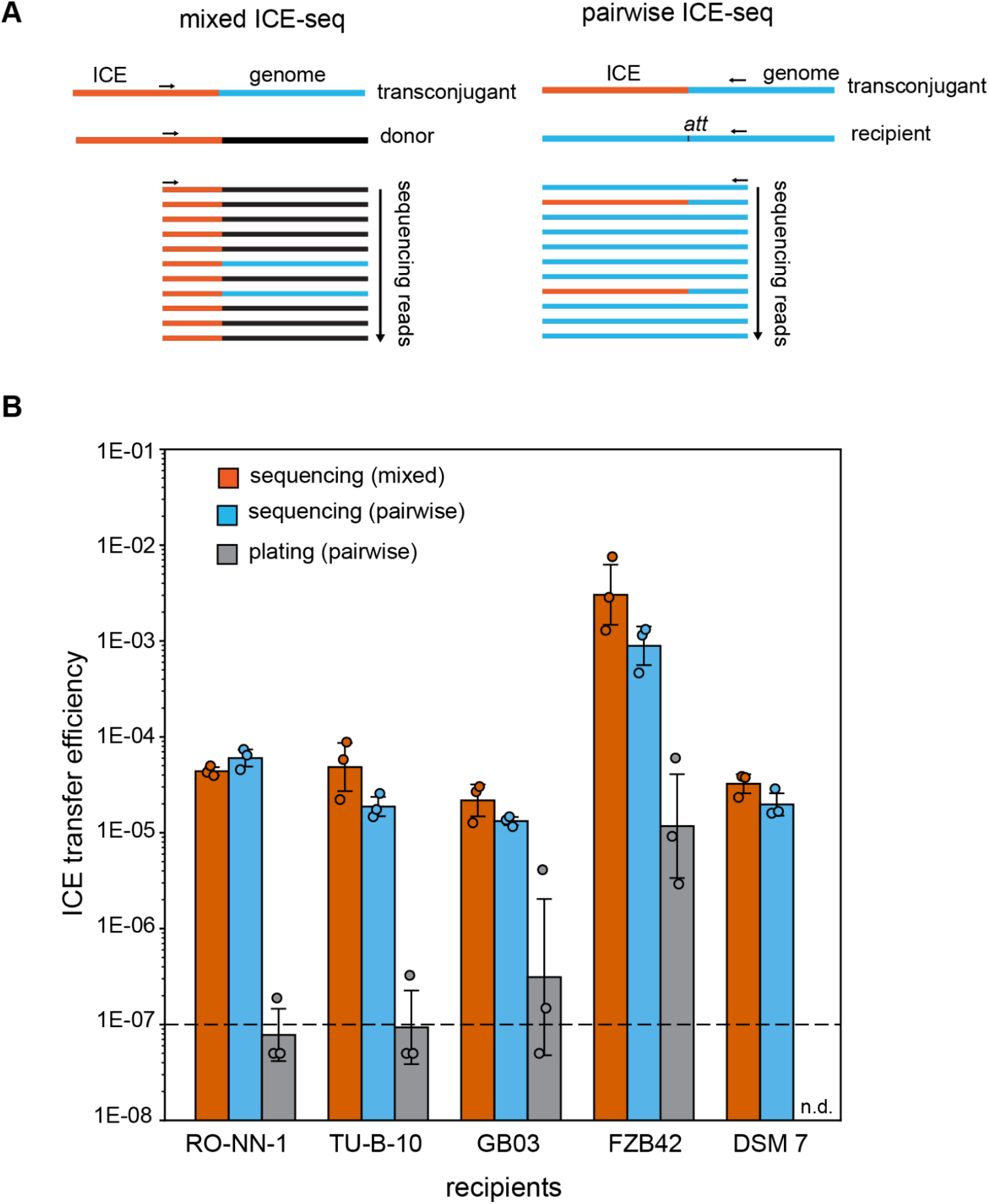
ICE-seq quantifies efficiency of ICE transfer into mixed *Bacillus* recipients. **A)** Primers in mixed assays amplify reads starting in the ICE and sequencing out into the surrounding genome, which allows for identification of any recipient. Primers in pairwise assays target the recipient genome to sequence into the *att* site. **B)** ICE matings were performed either with a mixture of five *Bacillus* recipients or in pairwise assays with each recipient individually.

The number of reads mapping to the donor, recipient, and transconjugant were quantified by ICE-seq. Transfer efficiency was calculated either by the number of transconjugant reads per donor reads in mixed assays, or by the number of transconjugant reads per reads per recipient reads in pairwise assays. Cells from the pairwise matings were also plated on selective plates and transfer efficiency was calculated as the ratio of transconjugant colony forming units (cfu) per recipient cfu. *B. amyloliquefaciens* strain DSM 7 did not grow with the minimal media needed for selection, so plating data were not included for this strain (n.d. = not determined). Error bars represent geometric mean ± 1 geometric standard deviation.

### RM systems play an important role in ICE transfer efficiency

In contrast to the quantitative similarity between the two sequencing assays, the phenotypic plating assays consistently measured lower ICE transfer efficiencies than sequencing (Figure 3B). We first hypothesized that this discrepancy was due to extracellular DNA from dead cells being detected by ICE-seq. However, using a DNase treatment to remove residual DNA prior to genome extraction did not alter the sequencing results (Fig. S2). We next hypothesized that ICE transfer frequently led to cell death but not lysis, possibly due to microbial defense systems. RM systems are known to restrict transformation by dsDNA, but are thought to have a minor effect on conjugative transfer of ssDNA such as ICEs (38, 47). To test this hypothesis, we experimentally identified methylation motifs in each of the recipients. Methylation motifs for *B. velezensis* GB03 and FZB42 were previously identified by nanopore sequencing (De Valle, in prep), and a motif for *Bacillus subtilis* subsp. *subtilis* RO-NN-1 was previously identified by PacBio sequencing (48). The methylation for *B. spizizenii* TU-B-10 was unknown, so we performed PacBio sequencing and identified 6-methyladenosine methylation at the motif ‘TGG**A**NNNNNNNTTG.’ Next, we calculated the occurrence of each motif in ICE*Bs*1 (Fig. S3). There was a strong log-linear correlation between the number of motifs in the ICE and the transfer efficiency as measured by plating (Fig. 4A). These results suggest that recipient restriction enzymes target these motifs when incorrectly methylated and cleave the ICE once transferred into the new cell. The logarithmic relationship between the number of methylation motifs and the transconjugant survival is consistent with independent cleavage of each motif.

**Figure 4.**
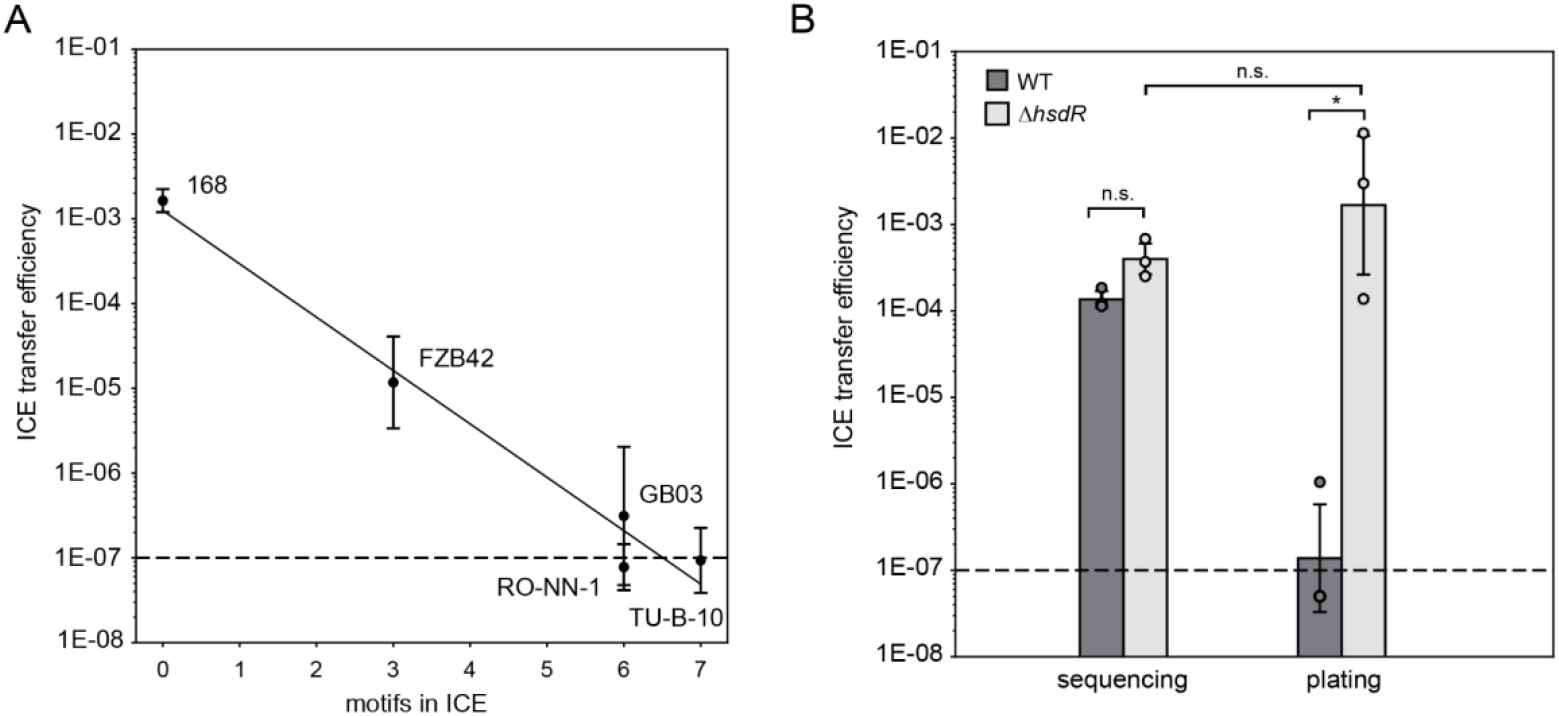
Restriction modification systems affect ICE transfer. **A)** ICE transfer efficiency by plating decreases as the number of restriction enzyme-targeted motifs in the ICE increase. **B)** A Type I restriction enzyme was deleted in the *Bacillus subtilis* subsp. *subtilis* RO-NN-1 recipient and conjugation efficiency was measured by ICE-seq and plating (efficiency = transconjugants/recipients). By sequencing, conjugation efficiencies of the mutant and wild type were not significantly different, while by plating the difference was significantly different (n=3, t-test, P<0.05). Error bars represent geometric mean ± 1 geometric standard deviation.

If RM systems were responsible for the difference between the sequencing and plating results, we hypothesized that deleting a restriction enzyme that targets the ICE from a recipient strain would not affect the sequencing results but would cause a large difference in the plating results. To test this hypothesis, we compared ICE transfer into *B. subtilis* RO-NN-1 and an otherwise isogenic strain with a deletion in the *hsdR* gene encoding a Type I restriction enzyme (48). The motif targeted by HsdR occurs six times in ICE*Bs*1 and is the only motif methylated in *B. subtilis* RO-NN-1. As predicted, results by sequencing in these two strains were statistically indistinguishable, while plating results varied by approximately three orders of magnitude (Figure 4B, n=3, t-test, P<0.05). Based on these results, we concluded that restriction enzymes in the recipient strain are likely targeting unmethylated motifs in the ICE, leading to double-stranded breaks in the genome and cell death (Fig. S4).

### ICE-seq tracks ICE transfer in a synthetic soil community

To test whether ICE-seq can measure ICE transfer in complex microbial communities like those found in the plant microbiome, the ICE donor was mated with a synthetic soil community composed of 150 phylogenetically-diverse bacteria isolated from poplar (Fig. 5A) (49). ICE-seq was used to track any potential ICE transfer to all members simultaneously. In this community there are nine strains that contain ICE*Bs*1 *att* sites (*Bacillus* sp. OV322, *Bacillus* sp. OV166 (has two *att* sites), *Bacillus* sp. YR288, *Bacillus* sp. BK245, *Bacillus* sp. BK100, *Psychrobacillus* sp. OK032, *Psychrobacillus* sp. OK028, *Bacillus* sp. OV752, and *Viridibacillus* sp. OK051). Unique kmers downstream of the *att* site in these strains were determined, and reads that matching these recipients were counted. Of these strains, ICE-seq detected only one recipient that obtained and integrated the ICE in its *att* site, *Bacillus* sp. BK100, which is closely related to *Bacillus velezensis* (Fig 5B). To validate this sequencing result, ICE transfer into BK100 was also measured in pairwise phenotypic assays, and transconjugant colonies were isolated and sequenced to confirm ICE integration (Fig. S5). Based on results for pairwise conjugations using purified isolates, ICE*Bs*1 has been proposed to transfer broadly within Gram-positive strains, including strains lacking the canonical *att* site (21). However, we observed ICE transfer and integration only in the clade of strains closely related to *B. subtilis* (Fig. S6), suggesting that ICE*Bs*1 may be more phylogenetically restricted, particularly for conjugation in mixed communities. Because ICE-seq can detect transconjugants which do not ultimately survive due to restriction, RM systems are unlikely to cause this limited transfer breadth. There are many possible factors which could limit transfer range at any of the steps involved, including but not limited to, mating pair formation, DNA translocation, complementary strand replication, and genome integration.

**Figure 5.**
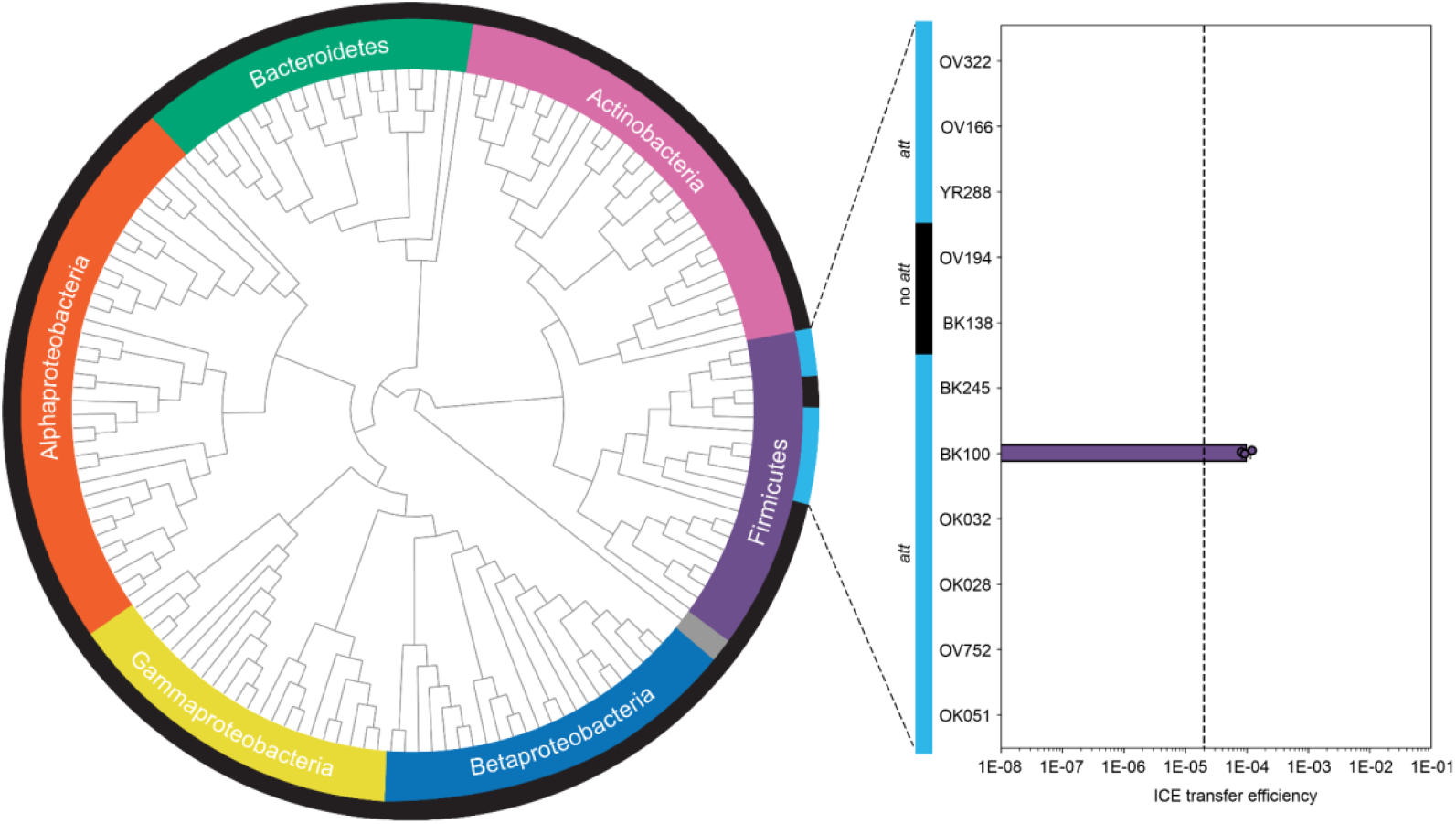
ICE transfer and integration is highly conserved. **A)** A phylogenetic tree was constructed of the 150-member poplar-isolated community. A cyanobacteria *Prochlorococcus marinus* (NR_125480) was used as the outgroup (grey) **B)** *Bacillus sp*. strain BK100 was the only recipient of the 150 poplar isolates detected by ICE-seq that was able to integrate the ICE in its canonical *att* site. ICE transfer efficiency was quantified by the number of transconjugant reads per number of donor reads. Error bars represent geometric mean ± 1 geometric standard deviation.

## Conclusions

In this work, we developed a high-throughput sequencing technique to track and quantify ICE transfer into multiple recipients simultaneously. Unexpectedly, validation of the ICE-seq method demonstrated that RM systems play an important role in limiting ICE transfer. HGT specificity is important for targeted microbiome engineering to prevent possible detrimental ecological off-target effects. Since RM systems vary widely even within species, altering ICE methylation may provide a way to target ICE transfer into individual bacterial strains within mixed communities. Furthermore, by tracking ICE transfer into 150 members of the poplar microbiome simultaneously, we determined that ICE*Bs*1 transfer in mixed communities is more restricted than expected. Additionally, because ICE-seq detects all ICE integration events, even those into recipients that ultimately do not survive, this technique can be used to capture the breadth of ICE transfer in complex microbiomes and identify recipients that other techniques may miss.

## Methods

### Strains and media

Strains used in this study are listed in Table S1. *Bacillus spizizenii* TU-B-10, *Bacillus subtilis* subsp. *subtilis* RO-NN-1, *Bacillus subtilis* subsp. *subtilis* JH642 substr. AG147, *Bacillus amyloliquefaciens* DSM7, *Bacillus velezensis* FZB42, and *Bacillus velezensis* GB03 were obtained from the Bacillus Genetic Stock Center (BGSC). ICE donor and recipient strains (ELC1560 and ELC1008) were provided by Dr. Alan Grossman. *Bacillus* strains were grown in LB at 37°C and 250 rpm. Antibiotics and other additions were used at the following concentrations: kanamycin (20 μg mL^-1^), spectinomycin (100 μg mL^-1^), chloramphenicol (10 μg mL^-1^), erythromycin (1 μg mL^-1^), lincomycin (12.5 μg mL^-1^), D-alanine (100 μg mL^-1^), tetracycline (200 ng mL^-1^), IPTG (1 mM), agar (1.5% w/v). Davis minimal media is composed of 1X phosphate-citrate buffer (per liter: 10.7 g K_2_HPO_4_, 6 g KH_2_PO_4_, 1 g sodium citrate adjusted to pH 7.5 with KOH), 0.2% glucose, 3 mM MgSO_4_, 0.02 mM C_6_H_8_FeNO_7_, 14.5 mM C_4_H_6_KNO_4_. Spizizen’s minimal salts (Spiz) contains 2 g (NH_4_)_2_SO_4_, 14 g K_2_HPO_4_, 60 g KH_2_PO_4_, 1 g sodium citrate, 2 g MgSO_4_ x 7H_2_O per liter.

### Strains and construction

*Bacillus* transformants were obtained by natural competence following standard protocols (50)The construct for ICE donor *comK*:ermR allele replacement was amplified using genomic DNA from ELC1008 and transformed into strain ELC1560. All primers used are listed in Table S1. Transformants were selected using LB medium and appropriate antibiotics. Engineered strains were verified by whole-genome resequencing.

### ICE-seq validation

Recipient strains containing ICE*Bs*1 were transformed by conjugations with donor strain JMB299 and isolated from a single colony using Davis minimal media supplemented with kanamycin. These strains were confirmed using whole genome sequencing. Liquid cultures of these strains and JMB299 were grown overnight with kanamycin and diluted 100-fold the next morning. Once cultures reached mid-logarithmic growth, they were washed once and resuspended in a potassium phosphate buffer (PBS) and serial diluted. Cells were mixed to appropriate dilution ratios and directly used for genomic DNA isolation. Genomic DNA was library prepped using ICE-seq method previously described, sequenced, and analyzed using our bioinformatic pipeline.

### ICE-seq

Genomic DNA was extracted using Qiagen Blood and Tissue kit according to the manufacturer’s instructions. DNA was quantified by Qubit 4 fluorometer (ThermoFischer Scientific, Waltham, Massachusetts) and 800-1000 ng per library was sheared using a Q800R3 Sonicator (Qsonica, Newtown, Connecticut) for 3 min with 15 s pulse and 15 s pause or Bioruptor Plus sonicator (Diagenode, Liege, Belgium) for 12 cycles 30 s pulse and 90 s pause. SPRI AMPure XP beads (Beckman Coulter) were used to select for fragments of 500 bp per manufacturer’s instructions. A NEBNext library prep kit (New England Biolabs, Ipswich, Massachusetts) was used for end repair, deoxyribosyladenine (dA)-tailing, and adaptor ligation, according to manufacturer’s instructions. Junction enrichment was performed using Q5 High-Fidelity 2x Master Mix (New England Biolabs, Ipswich, Massachusetts) and custom indexing and and region-specific primers with cycling conditions of 95°C for 5 minutes, then 25 cycles of 98°C for 30 seconds, 70°C for 15 seconds, 72°C for 15 seconds, and a final extension step of 72°C for 2 minutes. Two sequential rounds of AMPure XP bead purifications were performed as a final cleanup. Libraries were quantified with the Qubit 4 fluorometer and size confirmed using a Bioanalyzer DNA1000 chip (Agilent, Santa Clara, California). Libraries were sequenced using Illumina Novaseq or Nextseq system with paired-end 250-bp reads.

### Data analysis

Sequencing data was analyzed using a custom Python script, see Data Availability. To filter out non-specific and low-quality reads, an initial primer search was performed on R1 reads beginning after the first six bases, reflecting a buffer sequence in the P5 primer design. These R1 reads and their corresponding reverse-complemented R2 reads were searched for known recipient and donor sequences. Unique kmers (k=31) were found at fixed positions from input reference sequences, with a minimum pairwise Levenshtein distance of 3. The kmer nearest the beginning of the read was used for analysis, as kmer detection decreases further from the sequence start due to variable insert length and 3’ sequence quality of reads. Diagnostic kmers were identified and counted in the sequencing data and counts per reference sequence were saved to a matrix. These counts were used to calculate ICE transfer efficiency as a ratio of transconjugants per donor in mixed assays or transconjugants per recipient in pairwise assays.

### ICE matings

Strains for ICE matings were grown overnight in selective media and diluted 100-fold the next morning. When cultures reached an OD_600_ between 0.2 and 0.3, 1 mM IPTG was added to the donor strain to induce ICE conjugation. Strains were grown for another hour until early stationary phase was reached. Equal numbers of donor and recipient cells were mixed to achieve a total of 5 OD·mL. Mixed cells were collected by vacuum filtration with 0.2 μm pore size, 100 mL capacity sterile cellulose nitrate filters (Thermo Fisher Scientific, Waltham, MA) and washed with approximately 5 mL of LB to remove any antibiotics. Filters were incubated on Spiz agar plates at 37°C for 2 h and resuspended in liquid Spiz medium. Cells were either diluted in liquid Spiz medium for plating or collected for DNA extraction and sequencing by centrifugation at 10,000 rpm for 10 min at 22°C. For plating assays, cell suspensions were serially diluted and 100 μL of appropriate dilutions were plated on selective plates: recipient (Davis minimal agar) or transconjugant (Davis minimal agar with 20 μg/mL kanamycin). Colonies were counted after approximately 40 hours of incubation at 37°C.

### PacBio sequencing

A 5 mL culture of *B. spizizenii* TU-B-10 was grown overnight at 37°C. High molecular weight genomic DNA was extracted using a Monarch HMW DNA Extraction Kit (New England Biolabs, Ipswich, Massachusetts) and sequenced at Maryland Genomics at the University of Maryland Institute for Genomic Sciences.

### 150-member community mating and analysis

150 members were grown in R2A media in 96-well plates at 30°C and 250 rpm. Fast growing strains were grown for approximately 24 h, and slow-growing strains were grown for approximately 48 h. Individual strains were combined and OD_600_ of the mixture was measured. Donor strain was grown overnight in selective media and diluted 100-fold the next morning and induced with IPTG at early-log phase as was done for other matings. ICE matings proceeded as described for other strains, but combined cells were allowed to filter-mate overnight (∼15 h) before cells were collected for DNA extraction and ICE-seq.

## Acknowledgements

This work was supported as part of the Secure Ecosystem Engineering and Design Science Focus Area, funded by the U.S. Department of Energy, Office of Science, Biological and Environmental Research under Award Number ERKPA17. We would like to thank Dr. Alan Grossman for providing us the ICE donor strain ELC1560, which we modified and used in our ICE mating assays. We would also like to thank the University of Tennessee Genomics Core for sequencing our ICE-seq libraries and Leah Hochanadel for genome resequencing of engineered strains.

## Author contributions

E.B.G. conducted the majority of the experimental work and writing. R.D.K.: Computational pipeline development. D.M.K.: Sequencing library design and methods development. J.K.M.: Conceptualization, funding acquisition, supervision, writing.

## Data availability

Analysis code is available at https://github.com/ryandkuster/ice_element. Strains and plasmids are available on request, pursuant to a Material Transfer Agreement.

